# Integration of Biodynamic Imaging and RNA-seq classifies chemotherapy response in canine diffuse large B-cell lymphoma

**DOI:** 10.1101/2020.09.11.290353

**Authors:** Jonathan Fine, Sagar Utturkar, Deepika Dhawan, Phillip San Miguel, Gaurav Chopra, John Turek, David Nolte, Michael O. Childress, Nadia A. Lanman

**Author notes:** Correspondence: Nadia A. Lanman, Purdue University, 201 S. University Street, West Lafayette, IN 47907.; and Michael O. Childress, Purdue University, 625 Harrison Street, West Lafayette, IN 47907. Twitter handle: @NadiaAtallah1.

## Abstract

Diffuse large B-cell lymphoma (DLBCL) is a common, aggressive cancer of notorious genotypic and phenotypic heterogeneity. A major challenge is predicting response to drug treatment, which has typically been done using genomic tools alone with little success. A novel method that incorporates phenotypic profiling for predicting the effectiveness of therapy for individual patients is desperately needed. BioDynamic Imaging (BDI) is a technique for measuring time-dependent fluctuations in back-scattered light through living tumor tissues to identify critical changes in intracellular dynamics that are associated with phenotypic response to drugs. In this study, BDI and RNA sequencing (RNA-seq) data were collected on tumor samples from dogs with naturally occurring DLBCL, an animal model of increasingly recognized relevance to the human disease. BDI and RNA-seq data were combined to identify correlations between gene co-expression modules and linear combinations of biomarkers to provide biological mechanistic interpretations of BDI biomarkers. Using regularized multivariate logistic regression, we combined RNA-seq and BDI data to develop a novel model to accurately classify the clinical response of canine DLBCL to combination chemotherapy (i.e. CHOP). Our model incorporates data on the expression of 4 genes and 3 BDI-derived phenotypic biomarkers, capturing changes in transcription, microtubule related processes, and apoptosis. This pilot study suggests that the combination of multi-scale transcriptomic and phenotypic data can identify patients that respond to a given treatment *a priori* in a disease that has been difficult to treat. Our work provides an important framework for future development of strategies and treatments in precision cancer medicine.

## Introduction

Diffuse large B-cell lymphoma (DLBCL) is a common, aggressive form of non-Hodgkin lymphoma diagnosed in approximately 25,000 human patients each year, 1/3 of whom will die from the disease(1, 2). This cancer is characterized by molecular and biochemical heterogeneity that have confounded the use of targeted drugs to improve cure rates from conventional chemoimmunotherapy (3). The current standard of care chemotherapy regimen for DLBCL is a combination therapy that combines rituximab, cyclophosphamide, doxorubicin hydrochloride, vincristine sulfate (Oncovin^®^), and prednisone (i.e. R-CHOP). Chemotherapy often fails due to drug resistance (4), and no targeted therapy has been developed which significantly improves survival (5-7). Challenges in predicting treatment efficacy for individual patients, coupled with a lack of understanding in the development of R-CHOP resistance mechanisms, is a significant cause of this difficulty. The International Prognostic Index, the most commonly used prognostic tool, is based on simple clinical attributes (8, 9). While there are molecular prognostic schemes based on gene expression (10-14), there is no available method for accurately predicting response to chemotherapy regimens in individual patients with DLBCL. Thus far, genetic analysis alone is insufficient to predict the response of individual cases of DLBCL to drug therapy.

In spite of the importance of developing personalized therapy in DLBCL, progress has been slow (4). Locus heterogeneity, an outbred population, and poor clinical documentation lead to difficulties in human cancer gene mapping and predictive model development (15). While murine models have led to important breakthroughs in DLBCL research (16, 17), these models have limited translational application to precision cancer treatment (18, 19). Dogs with DLBCL have been proposed as a valuable model in which to develop novel personalized medicine strategies for humans with this cancer. Pet dogs develop DLBCL at a high rate, and the disease has similar clinical features to those seen in humans (15, 20). Pet dogs with DLBCL are treated with CHOP and exhibit heterogeneity in their response to CHOP chemotherapy as seen in humans’ receiving R-CHOP, making the dog model especially appropriate for informing precision medicine strategies to treat human DLBCL (20, 21).

BioDynamic Imaging (BDI) has been proposed as an accurate method to discriminate between DLBCL that will respond favorably versus unfavorably to treatment (22). BDI is an optical imaging technology that records phenotypic responses of fresh, living, three-dimensional tumor tissues to chemotherapeutic drugs in the *ex vivo* environment (5-7, 22-24). This phenotypic profiling occurs through the use of Doppler spectroscopy to characterize change in subcellular motion within tumor tissues following drug exposure. In a clinical setting, these motion-based responses are recorded from fresh *ex vivo* tissue biopsies with intact tumor microenvironments. These responses are then statistically associated with clinical outcomes such as objective tumor response or survival time. Although preliminary results show that BDI classifies chemosensitivity of naturally-occurring DLBCL in dogs (22), the relationship of BDI data to molecular processes underlying a tumor!s phenotypic drug response has yet to be defined.

We hypothesized that transcriptomic profiling could be correlated with BDI biomarkers, allowing us to characterize BDI biomarkers based on biological processes. Furthermore, we hypothesized that combined analysis of subcellular motion and gene expression patterns would enable the development of a machine learning model that could accurately classify DLBCL biopsies as sensitive or resistant to CHOP chemotherapy

In this study, we use intracellular dynamics data obtained from BDI and combine it with gene expression data from RNA sequencing (RNA-seq), defining the Doppler spectroscopic signatures recorded by BDI in terms of discrete biological processes. We also show that this integration creates an improved classifier of clinical chemotherapy response in canine DLBCL. Due to the success of machine learning applications across chemistry and biology (25-27), we explore multiple machine learning methodologies used to develop this classifier and discuss the implications of this model’s predictions for the biology of these tumors. The novelty of this work is that it combines RNA-seq with BioDynamic imaging and provides a biological mechanistic interpretation of BDI biomarkers. The majority of similar work in the literature uses a single analytical technique to predict a clinical result whereas our results suggest that multi-scale modeling approaches could offer an improved method for predicting the clinical response of human cancers to anticancer drugs.

## Material and Methods

### Assessment of clinical endpoints

Clinical management of study animals is detailed in Supplemental Methods. The primary clinical endpoints were objective response to chemotherapy and progression-free survival (PFS). The metric for assessing objective response was a caliper-based measurement measurements of a minimum of 1 and maximum of 5 peripheral lymph nodes. Objective response was classified according to established criteria (28) and is detailed in **Table 1**. Complete remission (CR) was defined as complete absence of detectable cancer following CHOP treatment. Partial remission (PR) was defined as >30% (but <100%) reduction in the sum of the longest diameters of up to 5 peripheral lymph nodes. Progressive disease (PD) was defined as >20% increase in the sum of the longest diameters of up to 5 peripheral lymph nodes, or the appearance of new lesions. Stable disease (SD) was defined as measurable tumor burden not meeting the criteria for PR or PD. Progression-free survival was defined as the time in days from initiation of chemotherapy to detection of PD or to death from any cause, whichever came first. All dogs enrolled in this study underwent incisional wedge biopsy or surgical extirpation of a peripheral lymph node at the time of study entry (i.e. prior to initiation of chemotherapy) to provide tissue for histopathologic confirmation of DLCBL. Portions of these biopsy samples were reserved and processed for BDI and additional tumor biopsy portions were frozen in liquid nitrogen or homogenized in TRIzol reagent (ThermoFisher) within 30 minutes of harvesting for collection of tumoral RNA.

**Table 1.** Details on the nineteen dogs evaluated in this study. Each patient is assigned a unique sample ID for pre- and post-chemotherapy and a tumor sample barcode to represent the dog both before and after treatment. Since the post treatment RNA-seq data is only used to identify additional genes and not used for the creation of any models, the Dog Identifier is used in all future figures and tables. PFS = progression-free survival, CR= complete remission, PR= partial remission.

| Sample Number | Sample Identifier | Clinical Outcome | Breed | Best Overall Response | PFS (days) |
| --- | --- | --- | --- | --- | --- |
| 1 | 12BDI | Sensitive | Mixed breed dog | CR | 232 |
| 2 | 100BDI | Sensitive | German shepherd | CR | 252 |
| 3 | 80BDI | Sensitive | Mixed breed dog | CR | 294 |
| 4 | RL84BD | Resistant | Great Dane | CR | 98 |
| 5 | 95BDI | Sensitive | Basset hound | CR | 244 |
| 6 | RL52CC | Sensitive | Bloodhound | CR | 267 |
| 7 | 75BDI | Sensitive | Rottweiler | CR | 327 |
| 8 | 42BDI | Resistant | Bernese mountain dog | CR | 84 |
| 9 | LY54CJ | Sensitive | Mixed breed dog | CR | 237 |
| 10 | LY11JD | Sensitive | Standard poodle | CR | 286 |
| 11 | LY16JW | Resistant | Mastiff | CR | 97 |
| 12 | RL10KM | Resistant | Rottweiler | CR | 89 |
| 13 | RL69KG | Sensitive | Golden Retriever | CR | 293 |
| 14 | LY99LB | Resistant | Mixed breed dog | PR | 43 |
| 15 | 43BDI | Sensitive | Mixed breed dog | CR | 315 |
| 16 | LY54SM | Sensitive | Mixed breed dog | CR | 1,018 |
| 17 | 37BDI | Resistant | Mixed breed dog | PR | 43 |
| 18 | 94BDI | Sensitive | Mixed breed dog | CR | 273 |
| 19 | RL02YB | Sensitive | Mixed breed dog | CR | 284 |

### RNA-seq and analysis

All solid tumor specimens were stored in liquid nitrogen, while RNA samples were stored at −80°C until the time of RNA extraction. RNA extraction was performed using the RNeasy kit (Qiagen). Poly A selection and library preparation was performed using the TruSeq Stranded kit (Illumina) followed by sequencing on a NovaSeq6000. Read trimming of the 2×150 reads was performed using Trimmomatic v.0.32 (29) followed by mapping to the CanFam3.1 genome using STAR v.2.5.4b (30) and counting using HTSeq v.0.7.0 (31). A differential expression analysis was performed using the DESeq2 (32) package and later edgeR (33-35) to select intersecting differentially expressed genes (DEGs). The Benjamini-Hochberg method(36) corrected for multiple testing and genes with FDR < 0.05 were denoted as significant. ClusterProfiler (37) was used to perform pathway and enrichment analyses. Enrichment and associations were deemed statistically significant at an adjusted p-value of 0.05.

Data are available at GEO accession number GSE156220.

### BDI

Complete details of the physics and optical engineering behind BDI can be found in (38, 39) and details of the application of BDI for drug screening can be found in (6, 23). The procedures for using BDI for measuring the response to chemotherapy, including sample handling and stabilization procedures, can be found in (22). Fresh tumor biopsy samples are processed into multiple small tissue samples of approximately 1mm^3^ volume, then immobilized in single wells of a 96-well plate before application of *ex vivo* drug treatments. Drug treatments consisted of a DMSO control, CHOP combination chemotherapeutic agent, as well as the individual components of CHOP in RPMI 1640 medium containing 0.1% DMSO. The CHOP therapy drugs were prepared containing doxorubicin (10µM), 4-hydroxycyclophosphamide (5µM), vincristine (60nM), and prednisolone (0.6µM). The longitudinal time course consists of 18 Loops over 12 hours: 6 Loops prior to drug through 12 Loops after drug. During the measurement of a sample in a single well 2000 digital holography (DH) camera frames are acquired of each sample at 25 fps. Each digital hologram is converted to the image domain through spatial 2D fast Fourier transforms (FFTs) to create a single image-domain section, termed an optical coherence image (OCI). A stack of 2000 image-domain sections (optical coherence images) constitute a time series for each pixel. The sample is masked from the background by the choice of an intensity threshold. The pixel-based time series is converted to a fluctuation power spectrum through temporal FFTs. The resulting pixel-based spectra are averaged into a single spectrum for a given sample on a given Loop. The baseline spectrum S(ω,0) is defined as the average of the last 4 Loops spectra prior to the treatment. A series of logarithmic fluctuation power spectra at successive times and subtracting the average baseline is used to generate the drug-response spectrograms, which display the shift in Doppler spectral content of the sample over the course of the assay.

### BDI analysis and summarization

The drug-response spectrogram is defined by Equation 1:

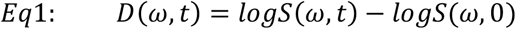

where the time index *t* represents the Loop number and *ω* represents the Doppler frequency. A spectrogram is generated for a given dog and a given treatment averaged over the wells. A data quality figure is assigned to each well based on multiple quality control criteria. The data quality has a base value of unity and is reduced by a factor of 2 for each criterion that is not satisfied by the well data. The spectrograms are averaged over the replicate wells for the treatment weighted by the data quality per well. Each dog’s response is thus represented by an average spectrogram for each treatment. This time-frequency representation next must be converted into feature vectors, which form the basis of BDI-derived biomarkers of a tissue’s phenotypic response to a given drug. The time-frequency spectrograms are converted into feature vectors with elements associated with parts or patterns of the spectrograms. In addition to spectrogram-based features, there are also preconditions (such as light-scattering brightness and spectral density dynamic range) as well as drug-induced changes in these preconditions.

All raw biomarkers are defined in **Table S1**. The time-frequency decomposition is approached globally and locally. Global patterns are generated as low-order Legendre polynomials. These polynomials are taken as an inner product over the spectrograms to generate Legendre coefficients that represent the global features of the spectrograms. Only orders 0, 1 and 2 are used along the frequency and time axes to generate 9 global features. Local patterns are simply low, mid, and high-frequency bands with average, linear and quadratic time dependence, which generates 9 local features. The preconditions consist of normalized standard deviation (NSD), backscatter brightness (BSB), number of pixels in the sample mask (NCNT), the spectral dynamic range (DR), the Nyquist floor (NY), the knee frequency (KNEE), the half-width (HW), the spectral slope (S) and the linear slope (SF). Each precondition is changed by the drug treatment, providing additional features that are the changes in the preconditions from baseline to the endpoint of the assay. Information on BDI biomarker definition is provided in the Supplemental Methods (**Tables S1-S3**). There are 27 drug-response features: 18 are based on spectrograms and 9 are drug-induced changes in preconditions, which are concatenated for each drug.

### Modeling of chemotherapy response

In early versions of our work, we investigated other machine learning models, including Support Vector Machine (SVM), Partial Least Squares (PLS), Random Forest (RF), and decision trees (DT). These models did not out-perform logistic regression, a finding congruent with other works which attempt to model clinical results using machine learning techniques(40). This can be explained by the fact that other machine learning models typically become over trained with such small training sets (a fact which we discovered early on in the modeling process). Additionally, these models are more difficult to analyze and perform feature selections upon, so given our early success with logistic regression we decided to not pursue additional models. Models were trained on data and feature selection was performed by selecting BDI biomarkers with greater than a 0.90 area under the precision recall curve (AUPRC) and high confidence protein coding DEGs, identified by both edgeR and DESeq2 as statistically significant and with greater than a 0.95 AUPRC. The Cohen Kappa statistic (41) was used to quantify model success based on inter-model reliability due to the unbalanced nature of the data. The resistant tumor samples were designated as the positive case and the sensitive tumor samples designated as the negative case. Kappa values of 0.80-0.90 are interpreted as strong, and above 0.90 is near perfect predictive ability (42, 43). The Caret package (44) was used to train models using leave-one-out cross validation (LOOCV) for hyperparameter tuning. Leave one out testing (LOOT) was used to enable testing of the final model.

### Cross-correlation of BDI and transcriptomic variables

A network analysis was performed, identifying modules of co-expressed genes using WGCNA (45). Log(FPKM) values were used after filtering genes with <0.3 FPKM in 20% of samples. A soft thresholding power of 14 was chosen, which was the lowest power at which the scale-free topology had a fit R^2^ of greater than 0.85 (46, 47). A signed adjacency matrix was computed and hierarchical clustering followed by a dynamic tree-cut algorithm identified modules with a minimum size of 30 genes. The first principal component of each of the modules (eigengenes), were used to correlate the modules with BDI biomarkers.

## Results

### RNA-seq and BDI capture transcriptional and phenotypic changes associated with chemotherapy resistance

Initially, we set out to characterize transcriptional and phenotypic changes associated with CHOP resistance in canine DLBCL. Importantly, the generation of RNA-seq data also provides an opportunity to map between gene expression and the BDI biomarkers, thus improving the interpretability of BDI results. RNA-seq and BDI were performed on tumors from nineteen pet dogs with DLBCL (**Table 1**). Of these tumors, 6 were categorized as resistant and 13 were considered sensitive to CHOP. Resistant tumors were considered those from dogs with a PFS of approximately ≤ 90 days. Previous work suggests that dogs with such tumors are in the lowest quartile for PFS (21), a subpopulation that derives negligible benefit from CHOP.

The canine RNA-seq data showed a high degree of variability between dogs (**Figure S1**), a characteristic also observed in human DLBCL (48). A total of 70 differentially expressed genes (DEGs) were identified between sensitive and resistant dogs, 27 of which are downregulated, and 43 are upregulated in resistant tumors (**Figure 1A, Table S4**). Gene expression changes mirror those identified in studies of R-CHOP resistance in human DLBCL patients. The most significantly enriched pathway amongst DEGs (**Table S5)** shows activation of the Jak-Stat pathway in the CHOP-resistant tumors (adjusted p-value=1.81×10^−4^), drawing a similarity between human and canine DLBCL (49). The gene set enrichment analysis (GSEA) (50) (**Figure 1B, Table S6**) likewise highlights a number of processes that are also involved in R-CHOP resistance in humans, while no enriched pathways were identified in sensitive samples (FDR<0.05). The oxidative phosphorylation pathway has previously been implicated in chemotherapy resistance, including in DLBCL (51, 52). Additionally, *MYC* targets are enriched in resistant samples, a finding that fits with the pathogenic role that constitutively activated MYC plays in human B-cell lymphomas (53). In addition to these pathways, the mTor signaling pathway, the complement and coagulation cascades, allograft refection, inflammatory related processes, and E2F targets were all shown to be enriched, similar to what has been shown in drug-resistant and -refractory DLBCL in humans (54).

**Figure 1.**
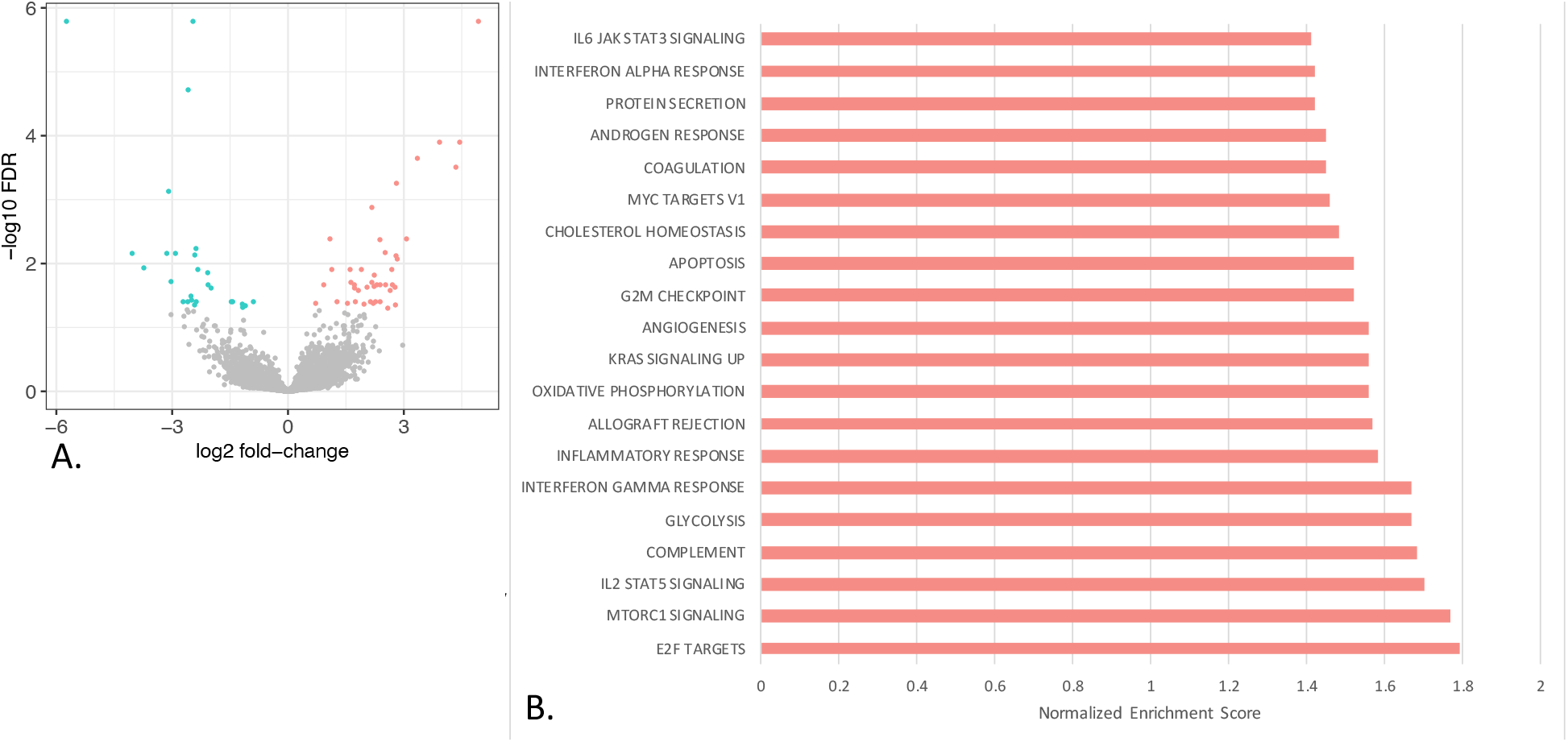
RNA-seq differential expression and enrichment results. **A**.Volcano plot with statistically significantly differentially expressed genes colored pink if they have higher expression in resistant samples and blue if they have higher expression in sensitive samples. **B**. Barchart showing the normalized enrichment score (NES) for the gene set enrichment analysis (GSEA) for significantly enriched gene sets at FDR<0.05. Positive enrichment scores indicate that the genes in the gene set have higher expression in resistant samples than sensitive samples. No genes sets were significantly enriched in sensitive samples.

BDI generated 81 biomarkers resulting from the 5 drug treatments (combination CHOP, along with individual monotherapy drugs), which are provided in **Table S7**. While these BDI time-frequency spectrograms provide robust measures of *ex vivo* changes in intracellular motion in response to CHOP combination and monotherapy treatment, the discrete biological mechanisms that underly each BDI biomarker were previously unknown, complicating biological interpretation of BDI data. For this reason, BDI and RNA-seq data were cross-correlated, in an attempt to bridge the gap between biological mechanism and BDI biomarkers.

### BDI biomarkers correlate with biological processes ranging from small to large-scale intracellular movements

Initially, a cross-correlation analysis was performed using a permutation test to quantify significance of correlations between BDI biomarkers and gene expression, followed by an enrichment analysis on the genes correlated with individual biomarkers. However, due to the large size of the data (81 BDI features and thousands of expressed genes), results were uninterpretable. To address this problem, genes and BDI markers were grouped into linear combinations of the individual features.

We had observed a number of significant linear relationships between biomarkers (**Figure 2A**), specifically in changes in preconditions as well as a few describing global changes across all frequencies and all times. Therefore, a principal component analysis (PCA) was used to generate a set of orthologous features for each drug treatment, providing us with sets of linear combinations of BDI biomarkers (**Table 2**). Within all drug treatments, the first 3 principal components contain the vast majority of the variability in the data, thus generating 15 features. The top 3 PCs contain 93.2% of the variability in the CHOP-, 88% in cyclophosphamide-, 79.3% in doxorubicin-, 90.6% in prednisone-, and 86.2% of the variability in vincristine-treated samples. Therefore, going forward, we focused on the 3 PCs that explain the majority of the variation in the data. For each of the five drug treatments, the loadings were used to determine the contribution of the original BDI biomarkers to each principal component.

**Figure 2.**
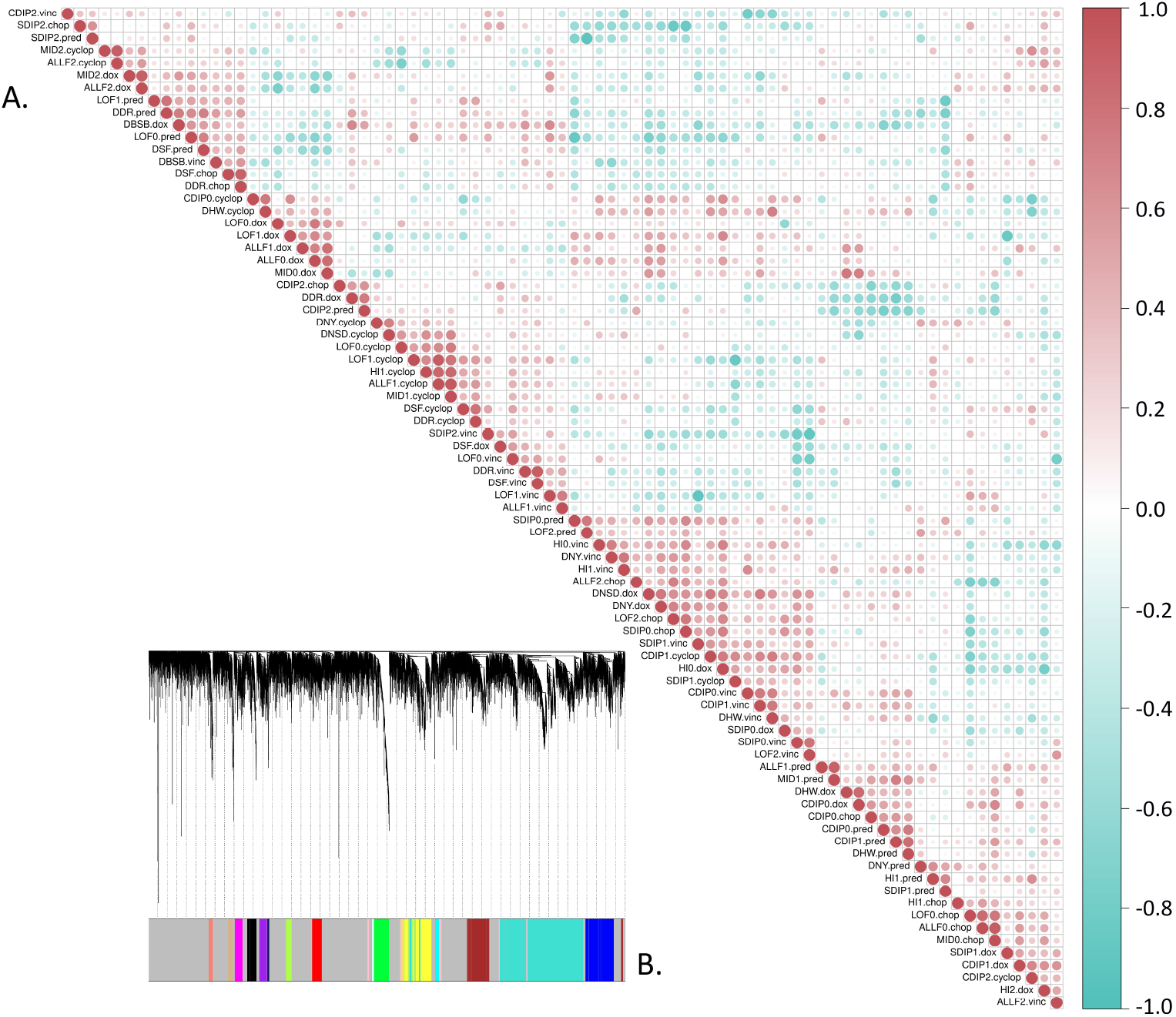
Linear relationships exist amongst BDI variables. **A**. A spearman correlation matrix shows linear relationships exist between many BDI variables. **B**. Gene dendrogram generated by average linkage hierarchical clustering on log transformed FPKM values. The color row underneath the dendrogram shows the module assignment determined by the Dynamic Tree Cut.

**Table 2.**
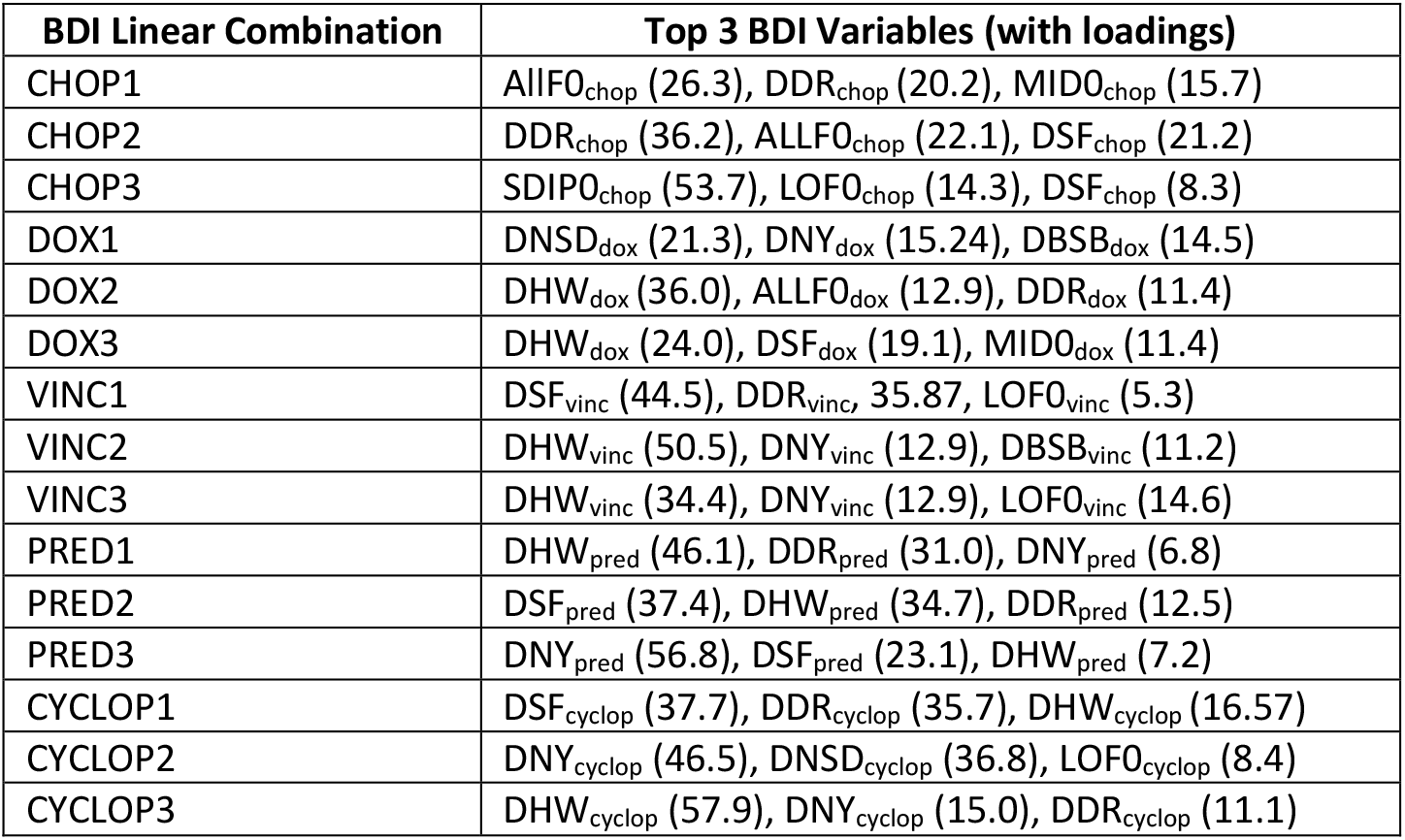
Table of the linear combinations of BDI biomarkers along with the major contributing raw biomarkers. BDI biomarkers are named according to the drug treatment followed by the principal component number. The top 3 raw BDI variables contributing the most to each linear orthologous BDI variable are shown, along with the loadings in parentheses.

We likewise summarized gene expression data into groups of correlated genes. Co-expression networks were identified in the RNA-seq data and correlated to BDI data. A total of 16 modules of co-expressed genes (**Figure 2B**), with sizes ranging from 44 to 2270 genes (**Table 3**) were identified. The networks were annotated with biological processes that are enriched in the genes grouped together in the modules (**Table S8**). Module eigengenes were then defined as the first principal component of the expression matrix of the corresponding co-expression module. The co-expression module eigengenes were cross-correlated with the top linear combinations of BDI biomarkers (**Table 3, Figure 3**) as well as the raw BDI biomarkers **(Table 3, Figure S2**). Here, we focus on correlation with modules that are enriched for processes that would result in changes in subcellular motion, which BDI detects and quantifies.

**Table 3.**
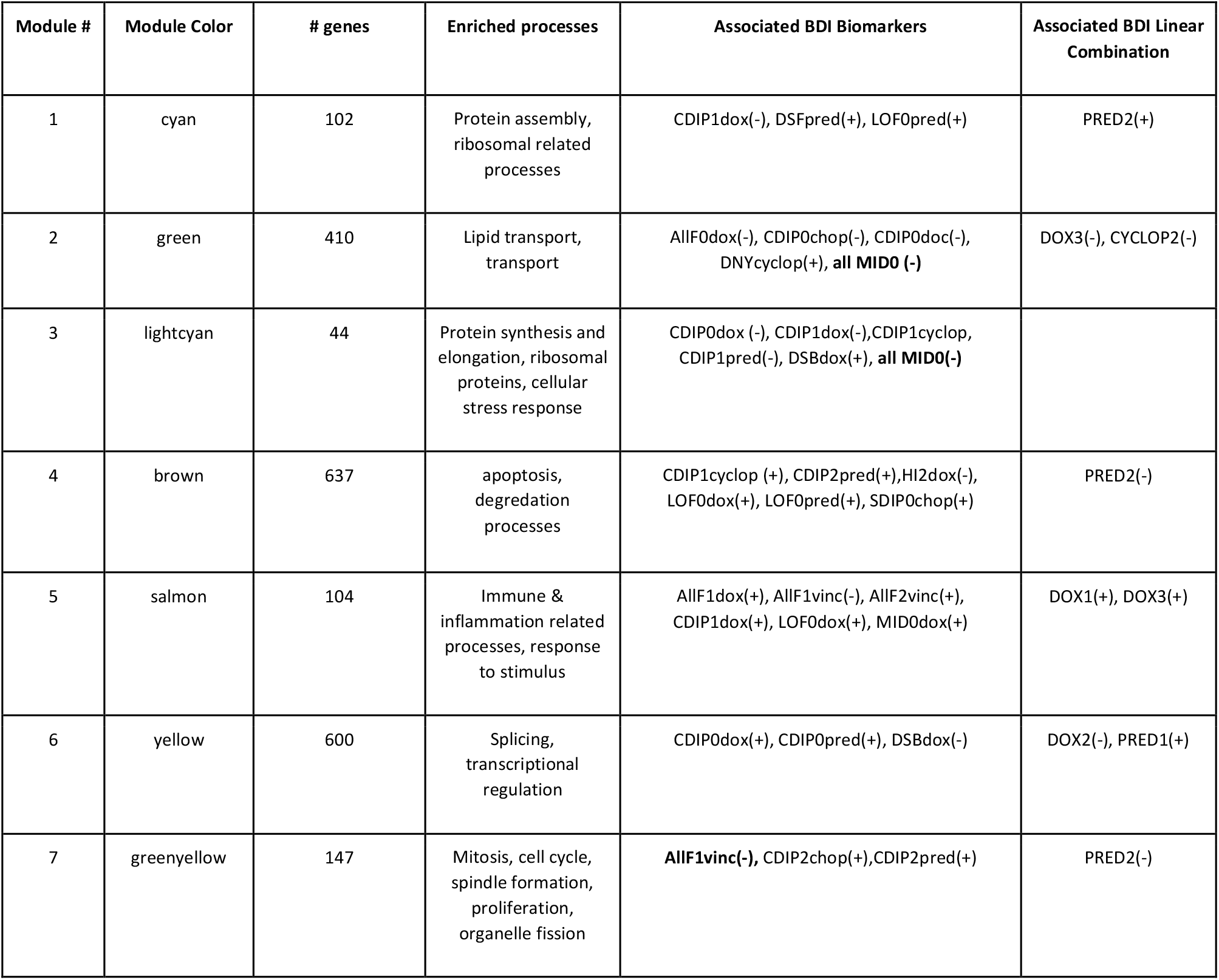
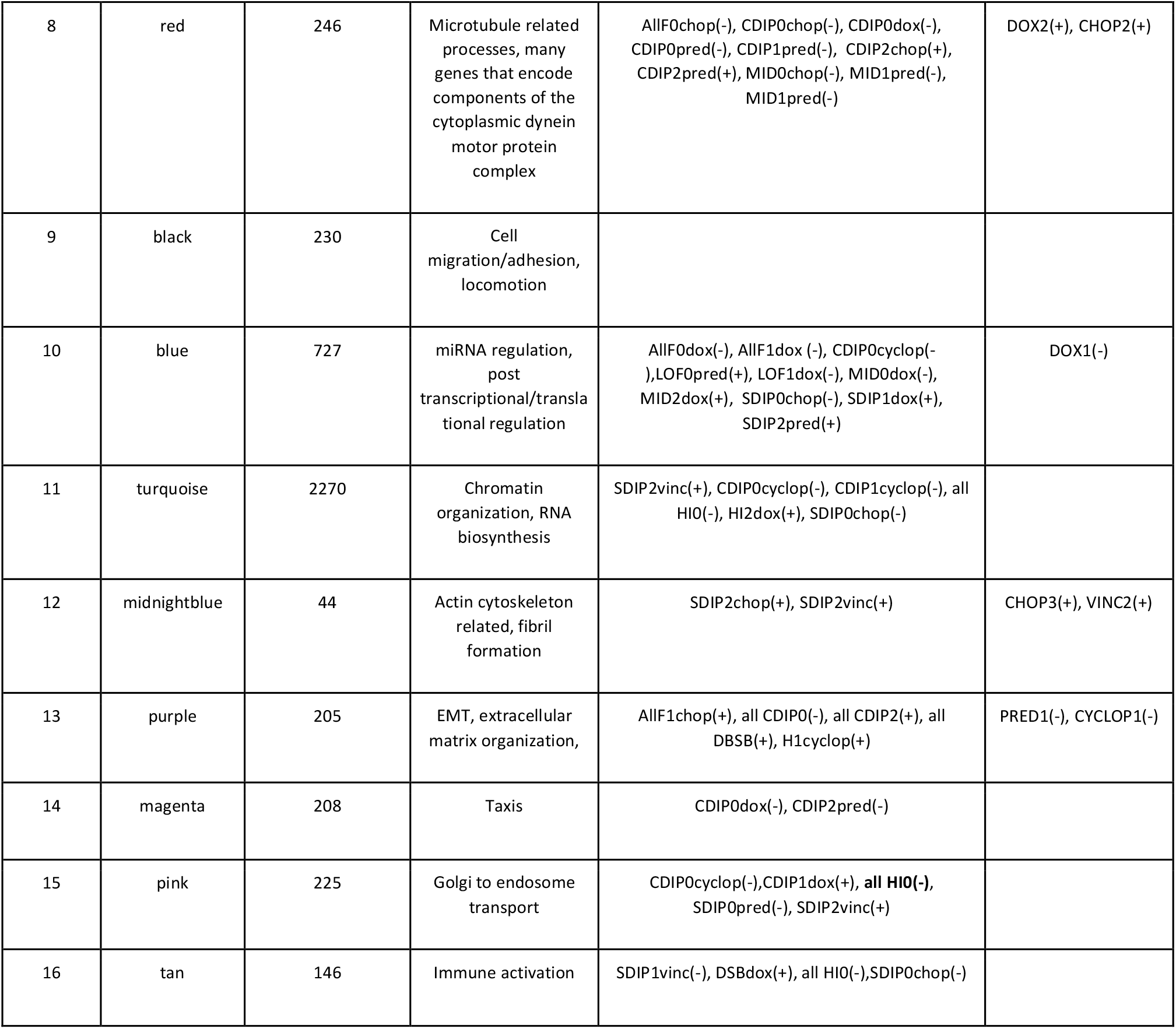
Modules of co-expressed genes, functions and associated BDI biomarkers. BDI biomarkers not correlated with other biomarkers were assessed individually for correlation with co-expression modules. The direction of the correlation (positive or negative) is shown in parentheses. The top positive and negative correlation (statistically significant and strongest linear relationship) for each BDI linear combination is shown.

**Figure 3.**
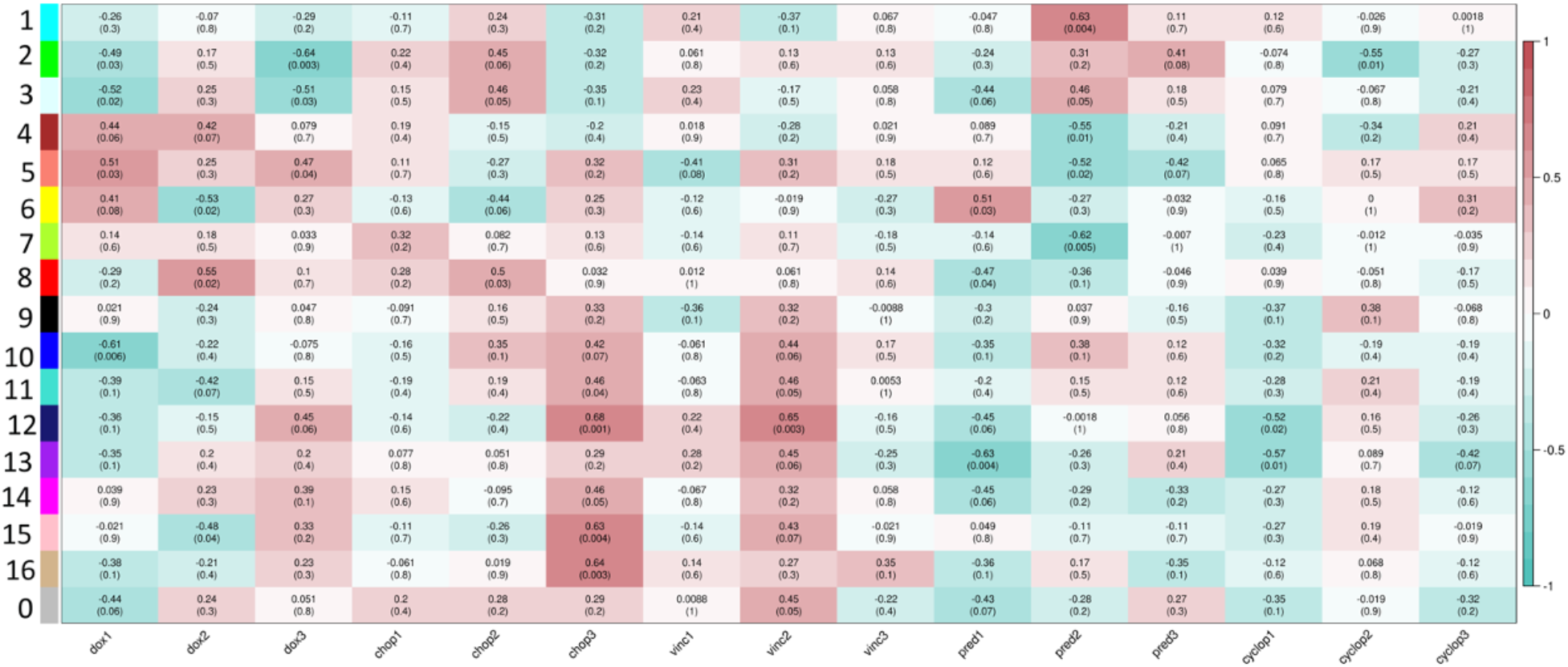
Network analysis and correlation with BDI biomarkers. Spearman correlations between identified co-expressed modules of genes and linear combinations of BDI biomarkers, generated by taking the first 3 principal components within each drug treatment. Colors indicate strength and direction of correlation and significance is indicated with adjusted p-values, provided in parentheses.

A number of precondition changes are related to large-scale intracellular movements and shape changes in cells through correlation of BDI linear combinations with co-expression modules 8, 12, and 13. Module 8 is strongly enriched for microtubule related processes including components of the dynein motor protein complex. Module 8 is correlated with changes in intracellular speeds (DHW) through linear combinations DOX2 and PRED1 (**Table 3**). Through the linear combination of CHOP2, the change in the dynamic range of spectral density (DDR) is likewise correlated with large-scale movement through Module 8. Changes in intracellular speeds (DHW) is correlated with modules 12 and 13, which correspond to changes in cellular shape through cytoskeleton and extracellular matrix reorganization, respectively(55) through VINC2 and PRED1. The change in the spectral slope (DSF) are likewise strongly correlated with changes in cellular shape (Modules 12 and 13, **Table 3**) through the linear combinations CHOP3 and CYCLOP1.

Correlations between co-expression modules and a number of raw BDI biomarkers with weak or nonexistent linear relationships with other markers were also observed. Golgi to endosome transport (Module 15, **Table 3**) is correlated with the global biomarker describing high frequency regions of the spectra across all times (HI0) under all drug treatments assayed. The correlation of the HI0 BDI biomarkers with a module enriched for Golgi to endosome transport fits with previous observations showing that high frequency regions of the spectrograms are indicative of fine movements, such as vesicle transport (24). The biomarker HI2_dox_, which captures quadratic time dependencies at high frequencies as well as LOF0_dox_ and LOF0_pred_, which captures low frequency regions of spectrograms across all times in doxorubicin and prednisone treated samples, respectively, are correlated with apoptosis (Module 4, **Table 3**) as are CDIP1cyclop and CDIP2_pred_. These correlations fit with previous experiments that correlated BDI markers to apoptosis through the use of drugs that induce mitochondrial toxicity and apoptosis (23).

All together, this cross-correlation analysis provides a set of testable hypotheses for novel interpretations of BDI biomarkers. This opens the door toward enhancing the utility of BDI and suggests that BDI can provide a means by which to draw biological conclusions about treatment effects in live *ex vivo* tissue biopsies. Likewise, these cross-correlation analyses also recapitulate known associations between BDI biomarkers and molecular processes, such as those previously observed using drug treatments known to alter vesicle transport and apoptosis (23, 24), providing increased confidence in the correlations between the biological functions enriched in co-expression modules and the BDI variables.

### Multi-scale data integration and modeling accurately classifies chemotherapy response

After BDI and RNA-seq data were collected from tumors, the goal was to determine whether a multi-scale model could be fit that would accurately classify tumor samples as sensitive or resistant to CHOP chemotherapy from pre-treatment data alone. The focus was on pre-treatment data alone in classifying samples because of the high impact such a model could have, in which a clinical outcome could be predicted before patients are needlessly subjected to chemotherapy that does not improve survival time. A total of 45 high confidence protein coding DEGs (**Table S9**) were combined with BDI biomarkers to yield a total input space of 126 features.

Traditionally, evaluating a machine learning model requires one to partition the data into three parts: a training set, a validation set, and a testing set. The training set is directly introduced into the machine learning model and the weights and other parameters of the model are identified from this set. This model is then applied to predict the results of the validation set and the model is allowed to be retrained multiple times using different hyperparameters to optimize performance. A final model is created using a combination of the training and validation sets. Note that the validation set is allowed to change multiple times so that it encompasses all of the data in the training set in a scheme referred to as *cross validation*, which allows the model hyperparameters to be adjusted for a large amount of data. The final model is then used to predict the results of the test set to obtain final statistics for the performance of the model. In addition to LOOCV, an evaluation paradigm called Leave One Out Testing (LOOT) was used to test models. Details of LOOT and a visual diagram of this process is given in **Figure S3**.

In these studies, we use absolute shrinkage and selection operator LASSO (L1) and ridge (L2) regularized logistic regression, which are established machine learning techniques that generalize well to data not seen in training or validation sets (40, 56). It should be noted that many of the BDI biomarkers were exceptionally strong predictors, and the three chosen were not the only such biomarkers identified. Rather, the low kappa value of 0.1894 in the model utilizing all 81 BDI biomarkers suggests that the model is overfit and thus aggressive feature selection methods were employed in the current setting.

To increase generalization, feature selection was employed based on variable Area-Under-the-Precision-Recall-Curve (AUPRC) (**Table S10-11**). BDI biomarkers with an AUPRC > 0.90 and high confidence RNA-seq variables with an AUPRC> 0.95 were selected. The 3 BDI biomarkers (SDIP1_dox_, LOF0_chop_, and ALLF1_pred_) yielded a model with a kappa of 0.43 and an accuracy of 73.68% (**Table S12**). The RNA variables selected were *ZFP92, KIAA1217, SH2D4A*, and *FGFR4* (**Figure 4A**). *ZFP92* encodes a zinc-finger protein 92 homolog (E-value= 1.66×10^−76^ in BlastP(57)), *SH2D4A* encodes a potential tumor-suppressor protein (58) and *KIAA1217* encodes a protein implicated in lung cancer progression (59). *FGFR4* encodes a receptor for fibroblast growth factors that is involved in cell proliferation, differentiation and migration(60) and is associated with poor prognosis in lymphoma (61) and in the development of drug resistance (62, 63). Two of the 3 BDI biomarkers (SDIP1_dox_, and LOF0_chop_) from the logistic regression model, were determined to be related to transcriptional/translational processes, and cellular shape changes through correlation with linear combination of BDI biomarkers CHOP3, respectively. Unfortunately, the BDI biomarker ALLF1_pred_ was not significantly correlated with any co-expression modules. It is possible that this biomarker will be defined through the analysis of a much larger dataset, which would no doubt increase the power of the cross-correlation analyses. There is no significant linear correlation between the RNA and the BDI biomarkers selected and a Principal Component Analysis shows that these 7 variables separate the 19 dogs (**Figure 4B-C**). A model trained on these 7 variables yielded a kappa of 1.00 with an accuracy of 100% (**Table 4, Table S13**), showing that the selected variables enable prediction of poor clinical response to CHOP therapy. Hence, combining gene expression and phenotypic data in a regularized multivariate regression model enables accurate classification of response to CHOP from pre-treatment data only, and therefore could significantly outperform models that incorporate data describing only gene expression(64) or phenotypic changes(22).

**Figure 4.**
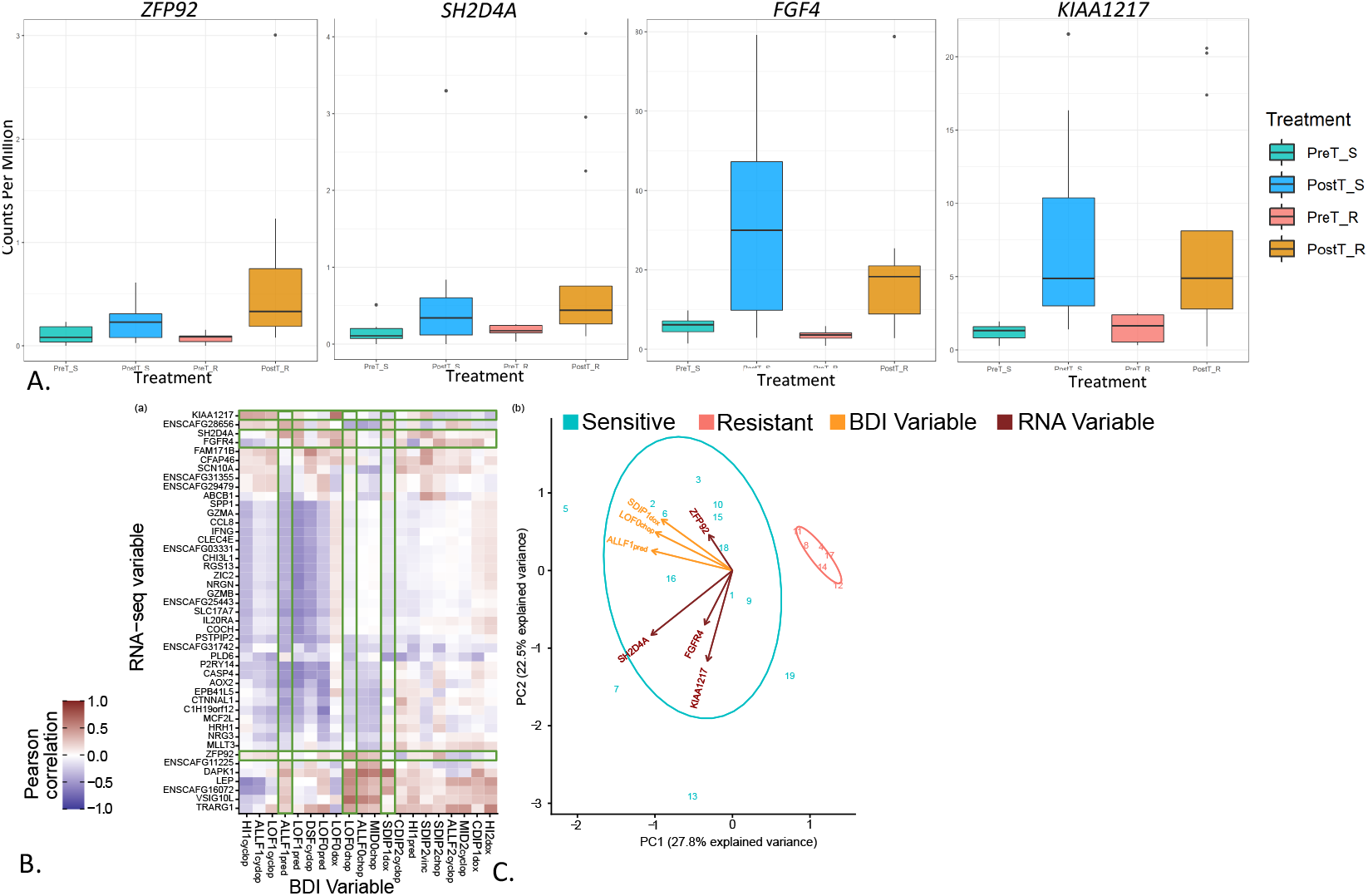
Visualization of logistic regression feature characteristics. **A**. Boxplots of expression of gene used in the model. **B**. Spearman Correlations between the top 20 BDI variables and RNA-seq variables where green boxes show the variables used to build the logistic regression model. **C**. A principal component analysis plot shows how the selected variables separate resistant versus sensitive lymphoma tumors.

**Table 4.** Summary of Leave One Out Testing. Top features are defined as the BDI and RNA variables with the highest AUPRC values.

| Model | Accuracy | Precision | Recall | F1 | TEST SET Kappa | MEAN Validation Kappa |
| --- | --- | --- | --- | --- | --- | --- |
| ALL BDI BIOMARKERS | 0.6316 | 0.4 | 0.3333 | 0.3636 | 0.1074 | 0.1894 |
| ALL RNA VARIABLES | 0.6842 | 0.5 | 0.1667 | 0.25 | 0.1094 | 0.3906 |
| TOP 3 BDI BIOMARKERS | 0.7368 | 0.57 | 0.6667 | 0.6154 | 0.4172 | 0.4272 |
| TOP 4 RNA VARIABLES | 0.8947 | 0.75 | 1 | 0.8571 | 0.7765 | 0.7671 |
| BDI AND RNA VARIABLES | 1 | 1 | 1 | 1 | 1 | 0.9922 |

## Discussion

This study illustrates the value of the canine model to study chemotherapy resistance to DLBCL and captures the response to therapy as well as the development of CHOP resistance in terms of both phenotypic and transcriptional responses. While a caveat of these studies is the small sample size, the study population was selected for clinical homogeneity. All dogs in this study had advanced-stage disease, yet none was affected by significant cancer-associated morbidity or comorbid disease. Thus, the DEGs and BDI biomarkers included in our models are likely to have captured biological variability inherent to the dogs’ cancers themselves rather than other host-related variables. The small sample size also resulted in our classifier “lumping” dogs with primary refractory DLBCL (two dogs with a best response of PR) together with those with initially responsive DLBCL that rapidly became CHOP-resistant (four additional dogs with CR lasting approximately 90 days; Table 1). It is probable that these two subpopulations of dogs have biologically and phenotypically distinct cancers, and that a classifier to resolve these two groups of dogs could be developed from a larger study population. Therefore, our results should be considered preliminary, and subject to revision as more data are collected.

We use cross-correlation of RNA-seq co-expression modules with BDI biomarkers to provide mechanistic interpretations of BDI biomarkers. These results both recapitulate previously known interpretations of biomarkers markers and identify novel interpretations of the BDI biomarkers. Such mechanistic interpretations of BDI biomarkers provides additional power to BDI, a technique that already has substantial clinical utility. In the future, these results should be confirmed both using a larger sample size as well as correlation with additional targeted drug treatments. A further improvement on the work performed herein would be to perform spatially resolved gene expression analyses to couple gene expression data with BDI data, which is already both temporally and spatially resolved. This would no doubt provide additional power for the identification of mechanistic interpretations of BDI data. Similarly, these analyses should be performed in additional samples to see whether these biological interpretations hold true in additional types of cancer as well as across other treatments.

We show that the integration of multiscale data describing cellular and molecular dynamics in a machine learning model classifies chemotherapeutic response in a relevant animal model of DLBCL. While the BDI biomarkers are strong predictors, the small sample size in this study necessitated aggressive feature selection. The three BDI biomarkers that we included in the most accurate models are not to be taken as the only important BDI biomarkers. Likewise, because different BDI biomarkers are associated with different biological processes, the BDI biomarkers with the most powerful predictive capacity may change depending on the treatment and cancer type assayed. Next, external validation should be performed on independent samples to validate the model. Future studies on targeted therapies as well as the cytotoxic drugs studied here should be performed with larger samples sizes.

We show that BDI is a powerful technology for predicting treatment response and provide biological interpretations of the BDI spectrograms. As this technology is a powerful method for characterizing cellular response to environments in living tissue, this resource is expected to aid in future interpretation of these data and the phenotypic responses accordingly. While genetic and transcriptomic profiles can be used to identify signatures for classifying samples, phenotypic variability often obfuscates the clinical validity of such methods. The studies here make a strong case for building predictive multivariate models that incorporate phenotypic and transcriptomic measures of drug response, such BDI, on live *ex vivo* tissue. Such approaches are needed in order to move personalized medicine forward and to predict therapeutic efficacy for individual patients.

## Acknowledgements

These studies were supported by the Walther Cancer Foundation, the Purdue University Center for Cancer Research (NIH grant P30 CA023168), the IU Simon Cancer Center (NIH grant P30 CA082709) and the Ralph W. and Grace M. Showalter Research Trust, and The Purdue University Challenge award. We gratefully acknowledge support of the Purdue University Genomics Core Facility, the Collaborative Core for Cancer Bioinformatics, and the staff at the Purdue University School of Veterinary Medicine. Additional funds in part by the Integrative Data Science Institute award at Purdue University and the National Institutes of Health, National Center for Advancing Translational Sciences ASPIRE Design Challenge awards are also acknowledged. We also greatly appreciate Purdue University Research Computing and Purdue ECN for continued computational and IT support. We are grateful to the clients at the Purdue University School of Veterinary Medicine, specifically those whom consented to have their beloved pets enrolled in this study.

## Authorship Contributions

Contribution: J.T., D.N., M.O.C., and N.A.L. conceived and designed experiments; J.F., P.S.M, J.T., D.N., M.O.C., and N.A.L. designed methodology; D.D, J.T, D.N., M.O.C. acquired the data, managed dogs, and provided facilities; J.F., S.U., G.C., D.N., M.O.C., and N.A.L. analyzed and interpreted the data J.F., D.N., S.U., and N.A.L wrote the paper and prepared figures; J.T., D.N., M.O.C., G.C, S.U., and N.A.L. reviewed/revised manuscript

## Conflicts of Interest Disclosures

David Nolte and John Turek have a financial interest in Animated Dynamics, Inc. that is seeking to commercialize biodynamic technology with a intellectual property license from Purdue University.

## Notes

### Summary of Updates

Figures are updated, with text enlarged for readability and extraneous figures placed into supplemental. Additional clinical details were provided and the manuscript was edited and reordered to improve readability.

