## Supplemental Information for "Integration of Biodynamic Imaging and RNA-seq classifies chemotherapy response in canine diffuse large B-cell lymphoma"

### **Supplemental Methods**

**Clinical Management of Study Animals**

Pet dogs with naturally occurring DLCBL were enrolled at the Purdue University Veterinary Teaching Hospital from August 2015-December 2016. Dogs were included in the study if they had histologically confirmed DLBCL affecting at least 1 peripheral lymph node that was > 2 cm in longest diameter, weighed >15 kg, and had expected survival time of at least 4 weeks with therapy. Dogs were excluded from the trial if they had received prior radiotherapy or chemotherapy of any kind (including glucocorticoids) or if they had high breed-associated risk for carrying the ABCB1-1Δ mutation affecting the MDR1 gene(1). Dogs with serious cancer-associated morbidity or evidence of other potentially life-limiting diseases also were excluded. The study protocol was approved by the Purdue Animal Care and Use Committee, and written informed consent obtained from each dog's owner prior to enrollment.

Seventy-five dogs were screened for enrollment in the trial, and 22 were enrolled. The reasons that dogs were not enrolled included: below minimum body weight requirement (n = 17), significant lymphoma-related illness or comorbid disease (n = 12), prior therapy for lymphoma (n = 7), suspected or confirmed non-DLBCL lymphoma (n = 7), owner declined (n = 6), high breed-associated risk for ABCB1-1Δ mutation (n = 3), and peripheral lymph nodes below minimum size requirement (n = 1). All enrolled dogs underwent standardized cancer staging tests at the time of study entry, which included complete blood count, serum biochemistry profile, thoracic and abdominal radiography, abdominal ultrasonography, and bone marrow aspirate cytology. A cancer stage was assigned to each dog based upon World Health Organization criteria(2).

All enrolled dogs were treated with the previously described chemotherapy protocol combining cyclophosphamide, doxorubicin, vincristine, and prednisone (i.e. CHOP,(3)). Tumor biopsy samples were scheduled to be collected from all enrolled dogs, both in the treatment-naïve setting and at the time cancer progression or relapse first became evident following initiation of CHOP. Three dogs were withdrawn from the trial at 5, 18, and 55 days after initiation of CHOP, respectively. The first two died at their owners’ homes of unknown causes (post-mortem exams were declined by the dogs’ owners). The third was withdrawn from the trial at the owner’s request after it experienced rupture of the right cranial cruciate ligament. Data on objective tumor response and progression-free survival from these three dogs were not submitted to statistical analysis, nor were RNA-seq data collected from any of these dogs’ pre-treatment biopsy samples. The remaining 19 dogs completed the trial and were evaluable for both objective tumor response and progression-free survival. The owners of 4 of these 19 dogs did not allow a second tumor biopsy to be performed at the time of cancer progression or relapse. The total dataset for analysis therefore comprised 34 (19 Pre-treatment and 15 post-treatment) tumor samples.

**Leave One Out Testing**

Leave One Out Testing (LOOT) is similar in spirit to Leave-One-Out Cross-Validation (LOOCV). LOOCV trains the model on all but one tumor (training set) and evaluates the resulting model on the tumor not used for training (validation set) to find the methodology and parameters which generalize best for the input features. Typically, these parameters are used to train a final model that is tested against an external dataset (test set) not used for LOOCV. In the current study, each sample was iteratively left out for LOOCV to enable testing of the final model on this ‘external validation set’ and this process was repeated in a similar manner to that of LOOCV. Thus in LOOT, LOOCV was iteratively performed after each sample was held out and in this manner we are able to be more stringent in validating and testing models than if LOOCV alone was performed. The cross-validated model obtained from the remaining observations is then used to predict the held-out member. This process is then repeated for all observations, allowing statistics to be calculated for the entire dataset that include the effects of hyperparameter optimization. In this methodology, the 19th sample is not used in training, and is therefore able to be used to measure this bias. The code required to perform analyses are at github.com/natallah/DLBCL_ml.

### **Supplemental Figures**

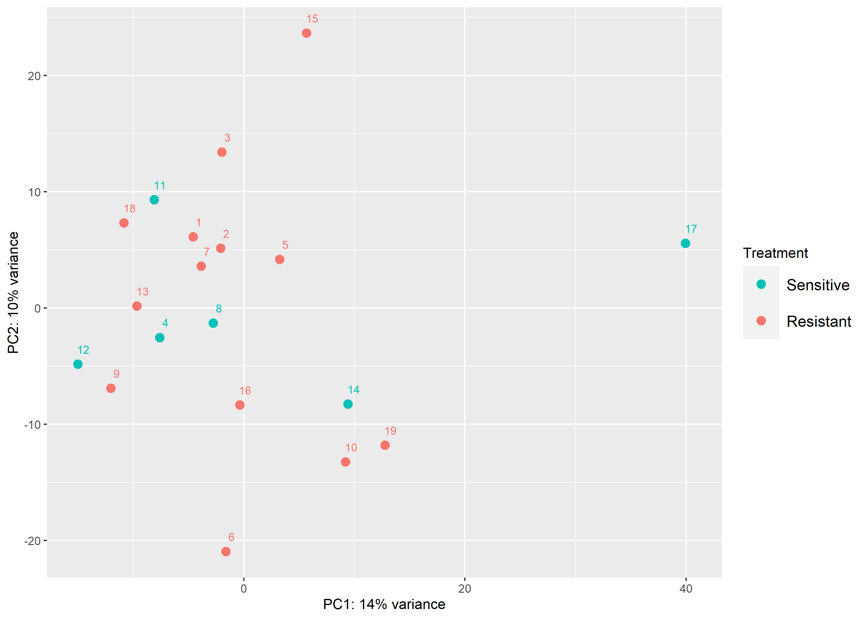

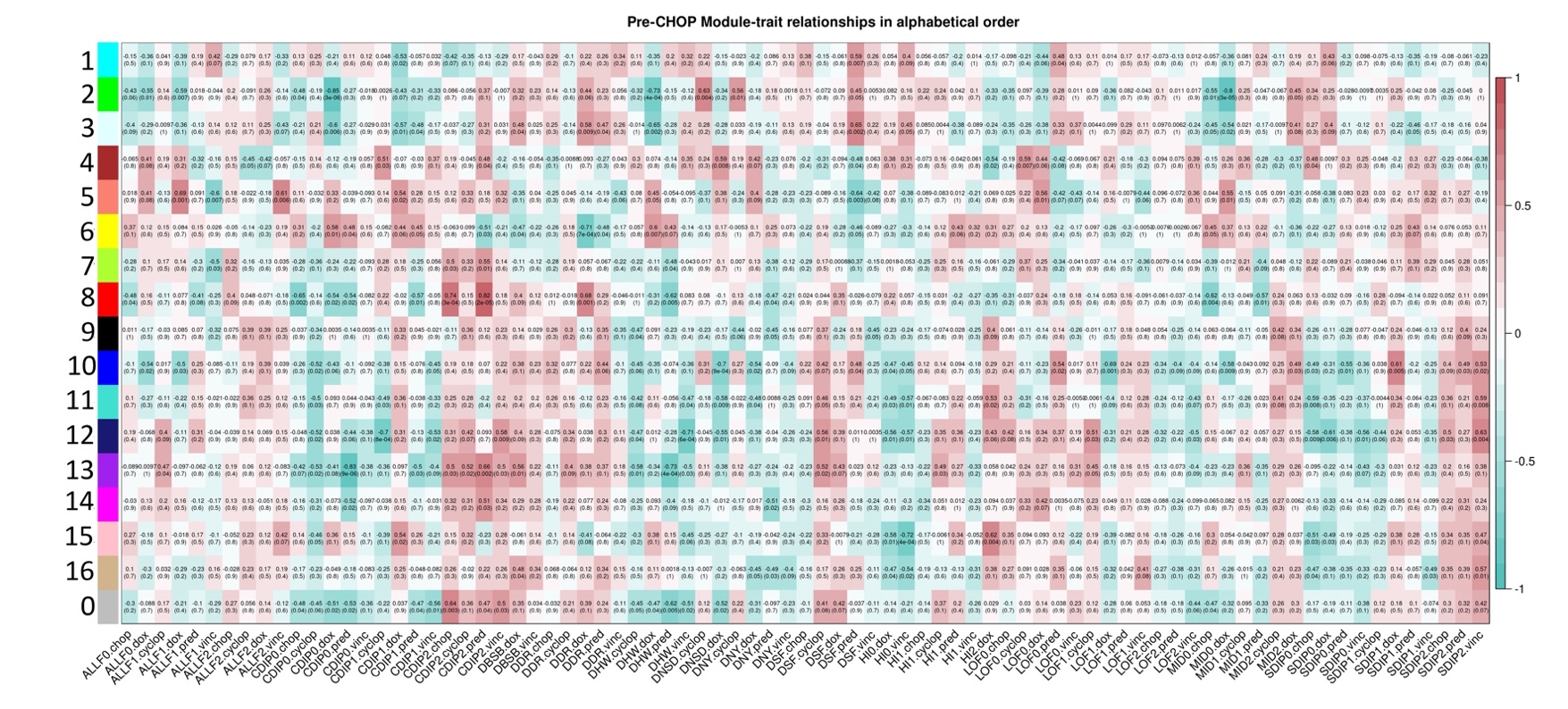
**Figure S1. A PCA plot of normalized variance stabilized RNA-seq data shows a high degree of variability between dogs.**

**Figure S2. Spearman correlations between identified modules and raw BDI biomarkers.**

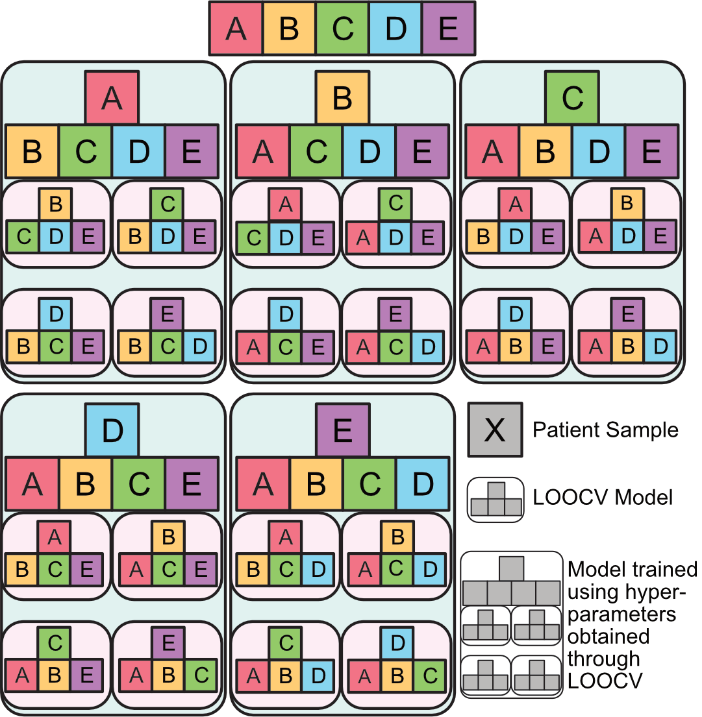

**Figure S3. A visual diagram of LOOT.** This figure shows the Leave-one-out testing (LOOT) process for 5 samples where A, B, C, D, and E are the observations. LOOT is similar in spirit to Leave One Out Cross-Validation (LOOCV) where a single observation is held out and the remaining observations are used to validate a potential model through LOOCV or other cross-validation methodology. The cross-validated model obtained from the remaining observations is then used to predict the held-out member. This process is then repeated for all observations, allowing statistics to be calculated for the entire dataset that include the effects of hyperparameter optimization.

### **Supplemental Tables**

**Table S1. BDI biomarker definitions.**

|  | **Biomarker** | **Description** |
| --- | --- | --- |
| **Global Spectral Biomarkers** | | |
| 1 | ALLF0 | All frequencies. All times |
| 2 | SDIP0 | Blue shift: All times |
| 3 | CDIP0 | Middle-out: All times |
| 4 | ALLF1 | All frequencies. Linear time dependence |
| 5 | SDIP1 | Blue shift: Linear time dependence |
| 6 | CDIP1 | Middle-out: Linear time dependence |
| 7 | ALLF2 | All frequencies. Quadratic time dependence |
| 8 | SDIP2 | Blue shift: Quadratic time dependence |
| 9 | CDIP2 | Middle-out: Quadratic time dependence |
| **Local Spectral Biomarkers** | | |
| 10 | LOF0 | Low-frequencies: All times |
| 11 | MID0 | Mid-frequencies: All times |
| 12 | HI0 | Hi-frequencies: All times |
| 13 | LOF1 | Low-frequencies: Linear time dependence |
| 14 | MID1 | Mid-frequencies: Linear time dependence |
| 15 | HI1 | Hi-frequencies: Linear time dependence |
| 16 | LOF2 | Low-frequencies: Quadratic time dependence |
| 17 | MID2 | Mid-frequencies: Quadratic time dependence |
| 18 | HI2 | Hi-frequencies: Quadratic time dependence |
| **Change in Precondition** | | |
| 19 | DNSD | Change in normalized standard deviation (NSD) |
| 20 | DBSB | Change in back-scatter brightness (BSB) |
| 21 | DNCNT | Change in number of pixels (NCNT) |
| 22 | DDR | Change in dynamic range (DR) |
| 23 | DNY | Change in Nyquist floor (NY) |
| 24 | DKNEE | Change in knee frequency (KNEE) |
| 25 | DHW | Change in half-width (HW) |
| 26 | DS | Change in Slope (S) |
| 27 | DSF | Change in linear slope (SF) |
| **Precondition** | | |
| 28 | NSD | Normalized standard deviation |
| 29 | BSB | Back-scatter brightness |
| 30 | NCNT | Number of pixels |
| 31 | DR | Dynamic range |
| 32 | NY | Nyquist floor |
| 33 | KNEE | Knee frequency |
| 34 | HW | Half-width |
| 35 | S | Slope |
| 36 | SF | Linear slope |
| 37 | B0 | Baseline: all frequencies |
| 38 | B1 | Baseline: linear frequency |
| 38 | B2 | Baseline: quadratic frequency |
| 40 | DQ | Data Quality |

There are approximately 5 spectral bands that can be defined in the drug-response spectrograms. These are defined in **Table S3** with their presumed biophysical origins.

**Table S2. Spectral Bands**

| **Band Name** | **Frequency Range** | **Speed Range** | **Biophysics Origins** |
| --- | --- | --- | --- |
| Cell Motility Band | 10 mHz | 3 nm/s | Crawling |
| Rheology Band | 12.5 mHz – 100 mHz | 4 nm/s – 30 nm/s | Shape change |
| Mid Band | 100 mHz – 1 Hz | 30 nm/s – 300 nm/s | Membrane/Nuclear |
| High Band | 1 Hz – 10 Hz | 300 nm/s – 3 µm/s | Organelle transport |
| Nyquist Band | 12.5 Hz | 4 µm/s | Vesicle transport |

Many drugs produce common spectrogram patterns. These are defined and described in **Table S4** along with presumed biophysical mechanisms.

**Table S3. Common Global Spectrogram Patterns.**

| **Pattern Name** | **Pattern** | **Characteristic Biomarker** | **Biophysical Origins** |
| --- | --- | --- | --- |
| Suppression | Blue | ALLF (neg) | Overall suppression of motion |
| Enhancement | Red | ALLF (pos) | Overall enhanced motion |
| Red Shift | Red - Blue | SDIP (neg) | Average speeds decrease |
| Blue Shift | Blue-Red | SDIP (pos) | Average speeds increase |
| Middle-Out | Red-Blue-Red | CDIP (pos) | Apoptosis/interrupted transport |
| Middle-In | Blue-Red-Blue | CDIP (neg) | More persistent transport |
| Drift Red | Time drift left | SDIP1 (pos) | Time drift to lower speeds |
| Drift Blue | Time drift right | SDIP1 (neg) | Time drift to higher speeds |
| Skew Red | Transition | CDIP1 and SDIP1(-) | Time change in pattern: red-shift |
| Skew Blue | Transition | CDIP1 and SDIP1(+) | Time change in pattern: bl-shift |

**Table S4. Differentially expressed genes between CHOP-sensitive vs -resistant dogs as calculated by DESeq2.** Genes were identified as differentially expressed if the adjusted p-values <0.05. Fold-changes were calculated as resistant/sensitive.

**Table S5. GO terms and pathways enriched amongst differentially expressed genes with adjusted p-value <0.05.**

**Table S6. Statistically significantly enriched Hallmark molecular signatures gene sets.** Gene sets were considered to be enriched if the FDR<0.05. No significantly enriched genes sets were identified in tumors that were sensitive to chemotherapy.

**Table S7. BDI biomarkers.** All 81 BDI biomarkers are available online at https://github.com/natallah/DLBCL_ml/blob/master/data/Supp_Tables_9_1_20 copy.xlsx

**Table S8. Enrichment results for modules of genes identified in WGCNA. Enriched pathways (KEGG and REACTOME), molecular signatures gene sets, and GO terms.** Full results with gene set enrichment included are available online at https://github.com/natallah/DLBCL_ml/blob/master/data/Supp_Tables_9_1_20 copy.xlsx

**Table S8. High confidence differentially expressed genes identified with an adjusted p-value <0.05 in both edgeR and DESeq2.**

**Table S9. The area under the receiver operator characteristic curve (AUROC) and area under the precision recall curve (AUPRC)** **for each of the BDI biomarkers.** This value is calculated where the ‘sensitive’ class is taken to be the control value and the ‘resistant’ class is taken to be the comparison value. These values are calculated independently of each other and represent how well each variable can be used to predict the clinical outcome. Here, an AUROC value of 0.5 would be obtained for a random classifier and values less than 0.5 indicates that an increase in the value corresponds to increased likelihood that the tumor is sensitive.

**Table S10. The area under the receiver operator characteristic curve (AUROC) and area under the precision recall curve (AUPRC)** **for each of the protein coding high confidence differentially expressed genes.** This value is calculated where the ‘sensitive’ class is taken to be the control value and the ‘resistant’ class is taken to be the comparison value. These values are calculated independently of each other and represent how well each variable can be used to predict the clinical outcome. Here, an AUROC value of 0.5 would be obtained for a random classifier and values less than 0.5 indicates that an increase in the value corresponds to increased likelihood that the tumor is sensitive.

**Table S11. Leave-one-out cross validation (LOOCV) results for regularized logistic regression models trained using various hyperparameters trained only using the BDI variables with at least a 0.80 area under the precision recall curve (AUPRC).** Here, L1 and L2 refer to Least Absolute Shrinkage and Selection Operator (LASSO) and Ridge regularization, respectively. The best kappa value is indicated with bold faced font.

**Table S12. Leave-one-out cross validation (LOOCV) results for regularized logistic regression models trained using various hyperparameters trained only using the high confidence protein coding differentially expressed genes with at least a 0.95 area under the precision recall curve (AUPRC).** Here, L1 and L2 refer to Least Absolute Shrinkage and Selection Operator (LASSO) and Ridge regularization, respectively. The models with the best kappa value are indicated with bold faced font.

**Table S13. Leave-one-out cross validation (LOOCV) results for regularized logistic regression models trained using various hyperparameters trained only using the high confidence protein coding differentially expressed genes with at least a 0.95 area under the precision recall curve (AUPRC) and with BDI variables with at least a 0.80 AUPRC.** Here, L1 and L2 refer to Least Absolute Shrinkage and Selection Operator (LASSO) and Ridge regularization, respectively. The best kappa value is indicated with bold faced font.
